# Diversification by CofC and control by CofD govern biosynthesis and evolution of coenzyme F_420_ and its derivative 3PG-F_420_

**DOI:** 10.1101/2021.08.11.456035

**Authors:** Mahmudul Hasan, Sabrina Schulze, Leona Berndt, Gottfried J. Palm, Daniel Braga, Ingrid Richter, Daniel Last, Michael Lammers, Gerald Lackner

## Abstract

Coenzyme F_420_ is a microbial redox cofactor that is increasingly used for biocatalytic applications. Recently, diversified biosynthetic routes to F_420_ and the discovery of a derivative, 3PG-F_420_, were reported. 3PG-F_420_ is formed via activation of 3-phospho-D-glycerate (3-PG) by CofC, but the structural basis of substrate binding, its evolution, as well as the role of CofD in substrate selection remained elusive.

Here, we present a crystal structure of the 3-PG-activating CofC from *Mycetohabitans* sp. B3 and define amino acids governing substrate specificity. Site-directed mutagenesis enabled bidirectional switching of specificity and thereby revealed the short evolutionary trajectory to 3PG-F_420_ formation. Furthermore, CofC stabilized its product, thus confirming the structure of the unstable molecule, revealing its binding mode and suggesting a substrate channeling mechanism to CofD. The latter enzyme was shown to significantly contribute to the selection of related intermediates to control the specificity of the combined biosynthetic CofC/D step. Taken together, this work closes important knowledge gaps and opens up perspectives for the discovery, enhanced biotechnological production, and engineering of coenzyme F_420_ derivatives in the future.

**Importance:** The microbial cofactor F_420_ is crucial for processes like methanogenesis, antibiotics biosynthesis, drug resistance, and biocatalysis. Recently, a novel derivative of F_420_ (3PG-F_420_) was discovered, enabling the production and use of F_420_ in heterologous hosts.

By analyzing the crystal structure of a CofC homolog whose substrate choice leads to formation of 3PG-F_420_, we defined amino acid residues governing the special substrate selectivity. A diagnostic residue enabled reprogramming of the substrate specificity, thus mimicking the evolution of the novel cofactor derivative and successfully guiding the identification of further 3-PG-activating enzymes.

Furthermore, a labile reaction product of CofC was revealed that has not been directly detected so far and CofD was shown to provide as another layer of specificity of the combined CofC/D reaction, thus controlling the initial substrate choice of CofC. The latter finding resolves a current debate in the literature about the starting point of F_420_ biosynthesis in various organisms.

## Introduction

Organic cofactors are small molecules other than amino acids that are required for the catalytic activity of enzymes (1). They are therefore crucial for the understanding of the biochemical and physiological processes their dependent enzymes are involved in, as well as the biotechnological exploitation of these enzymes. Coenzyme F_420_ is a specialized redox cofactor that was so far mainly identified in archaea and some actinobacteria (2). In archaea, F_420_ is a key coenzyme of methanogenesis (3). In mycobacteria, F_420_ plays a vital role in respiration (4,5), cell wall biosynthesis (6,7), as well as the activation of medicinally relevant antimycobacterial (pro-)drugs. For instance, the novel anti-tubercular drug pretomanid is activated by Ddn, an F_420_-dependent nitroreductase (8,9). In streptomycetes, F_420_H_2_ is used for reduction steps during the biosynthesis of antibiotics like thiopeptins (10), lanthipeptides (11), or oxytetracycline (12,13). Increasing interest in F_420_ is also driven by the utilization of F_420_H_2_-dependent reductases in biocatalysis, e.g., for asymmetric ene reductions (14-18).

Intriguingly, F_420_ also occurs in a few Gram-negative bacteria where it has been acquired most likely by horizontal transfer of its biosynthetic genes from actinobacteria (19,20). Initial studies have revealed that F_420_ is indeed produced by some of these organisms but their physiological role remains unknown (20,21). We have recently identified F_420_ biosynthetic genes in the genome of *Mycetohabitans* (synonym: *Paraburkholderia) rhizoxinica* (22), a symbiont that inhabits the hyphae and spores of the phytopathogenic mold *Rhizopus microsporus* (23-26). Surprisingly, we discovered that the symbiont produced a novel derivative of F_420_, which we termed 3PG-F_420_ (22). The cofactor activity of 3PG-F_420_ was comparable to classical F_420_ and could serve as a substitute for the latter in biocatalysis (22). Although this congener has not been described in any other organism, it could also be detected in the microbiota of a biogas production plant, thus demonstrating that it is not restricted to endofungal bacteria (22). The producers of 3PG-F_420_ in these habitats, however, are unknown.

The biosynthesis of 3PG-F_420_ (**Figure 1**) is generally similar to the biosynthesis of classical F_420_ (27). The pathway starts with the formation of the redox-active core moiety 7,8-didemethyl-8-hydroxy-5-deazariboflavin (F_O_) from L-tyrosine and 5-amino-6-ribitylamino-uracil, a reactive metabolite of the flavin biosynthesis pathway. The F_O_ core is then elongated by a chemical group that can formally be described as 2-phospho-L-lactate (2-PL) before an oligoglutamate tail is added. The biosynthesis of the 2-PL moiety has been the subject of several studies. Seminal work on archaea suggested that it is directly formed from 2-phospho-L-lactate: Incubation of cell extracts of *Methanosarcina thermophila* or *Methanocaldococcus jannaschii* with F_O_, 2-PL, and GTP led to the formation of F_420_-0 (28). Biochemical assays with purified CofC and CofD finally corroborated the model that the guanylyltransferase CofC catalyzes the reaction of 2-PL and GTP to lactyl-2-phospho-guanosine (LPPG) (29), which is then passed on to CofD to transfer the activated 2-PL moiety onto the precursor F_O_. However, the unstable nature of LPPG has prevented confirmation of its structure by NMR or mass spectrometry so far. The last biosynthetic step leading to the mature coenzyme F_420_ is catalyzed by the F_420_:glutamyl ligase CofE (30), which is responsible for the addition of the *γ*-linked oligoglutamate moiety to the F_420_-0 core, thus forming F_420_-n, with n indicating the number of glutamate residues.

**Figure 1:**
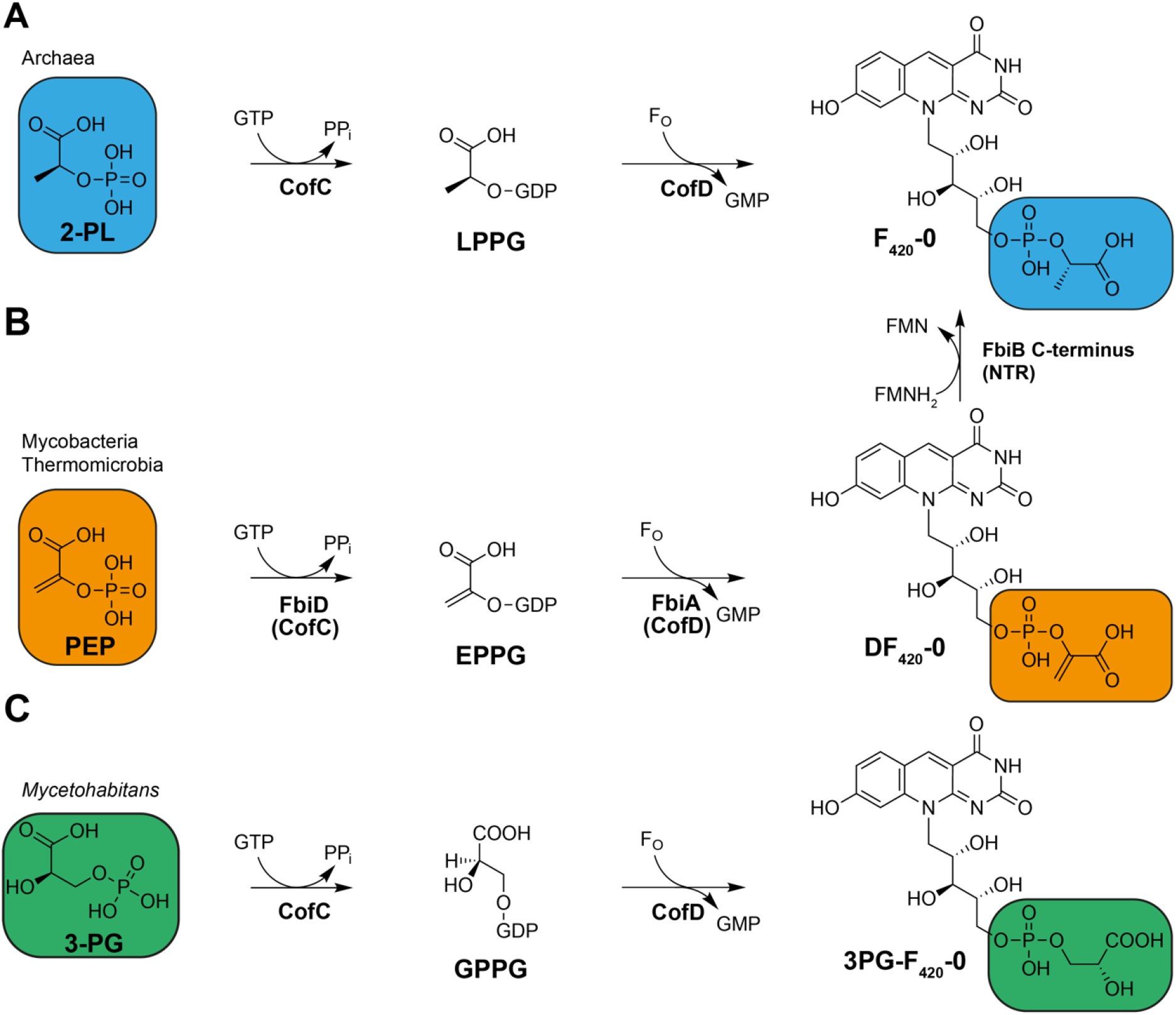
Biosynthetic steps performed by CofC and CofD during the formation of F_420_-species. **(A)** Formation of F_420_-0 from 2-PL as proposed for archaea. **(B)** Formation of the F_420_ precursor DF_420_, a pathway intermediate of F_420_ found in mycobacteria and Thermomicrobia. DF_420_ is further reduced to F_420_ by a nitroreductase (NTR)-like enzyme **(C)** Biosynthesis of 3PG-F_420_-0 in *M. rhizoxinica* and related endofungal bacteria. 2-PL: 2-phospho-L-lactate, 3-PG: 3-phospho-D-glycerate, EPPG: enolpyruvyl-2-diphospho-5′-guanosine, FMN: flavin mononucleotide, F_O_: 7,8-didemethyl-8-hydroxy-5-deazariboflavin, GMP: guanosine 5’-monophosphate, GPPG: 3-(guanosine-5’-diphospho)-D-glycerate, GTP: guanosine 5’-triphosphate, LPPG: L-lactyl-2-diphospho-5′-guanosine, NTR: nitroreductase, PEP: phosphoenolpyruvate, PP_i_: pyrophosphate.

In mycobacteria, CofE is not a free-standing enzyme but constitutes the *N*-terminal domain of the FbiB protein (31). It was shown recently that mycobacteria utilize phosphoenolpyruvate (PEP), but not 2-PL, to form F_420_-0. Instead of LPPG, EPPG is formed, which is converted into dehydro-F_420_-0 (DF_420_-0) by the action of FbiA, the mycobacterial CofD homolog. DF_420_-0 is then reduced to classical F_420_-0 by the *C*-terminal domain of FbiB, which belongs to the nitroreductase superfamily (32). We have shown that a similar pathway is present in the thermophilic bacterium *Thermomicrobium roseum* and related species (33). The formation of 3PG-F_420_-0, however, does not require any reduction step. Instead, enzyme assays revealed that 3-phospho-D-glycerate (3-PG) is activated by CofC, presumably forming the short-lived intermediate 3-(guanosine-5’-diphospho)-D-glycerate (GPPG), which is further transferred to the F_O_ core by the action of CofD.

However, it remained elusive, which amino acid residues within the CofC protein conferred the specificity switch towards 3-PG and how genetic mutation might have led to the evolution of 3PG-F_420_ biosynthesis. Furthermore, the question persisted, why the CofC CofD reaction only proceeds as a combined reaction and how reactive intermediates like LPPG are stabilized. Another open question concerned the role of 2-PL in the biosynthesis of F_420_ in archaea. While our previous data (22) matched seminal observations (29) of a substantial turnover of 2-PL by CofC enzymes of archaeal origin, other studies raised doubts that 2-PL is a genuine substrate of archaeal CofC homologs (32).

Here, we present a crystal structure of the 3-PG activating CofC from *Mycetohabitans* sp. B3 and revealed the amino acid residues governing 3-PG activation. By site-directed mutagenesis, we shed light on the evolution of 3PG-F_420_. We furthermore bring to attention that CofC strongly binds its product GPPG and collaborates closely with its partner CofD to control the flux of intermediates into the F_420_ biosynthesis pathway.

## Results

### Assessment of substrate specificities of CofC enzymes from several organisms

To gain a better understanding of CofC substrate specificities, we set out to identify more homologs of CofC accepting 3-PG as a substrate. We reasoned that related bacteria, harboring CofC homologs highly similar to the *M. rhizoxinica* enzyme (*Mrhiz*-CofC), would have a similar substrate preference. Indeed, using LC-MS we detected 3PG-F_420_ (**Figure 2A/B**) in cell extracts of *Mycetohabitans* sp. B3, a close relative of *M. rhizoxinica* that shares the same lifestyle as a symbiont of a phytopathogenic *Rhizopus microsporus* strain. Neither classical F_420_ nor DF_420_ was detectable.

**Figure 2:**
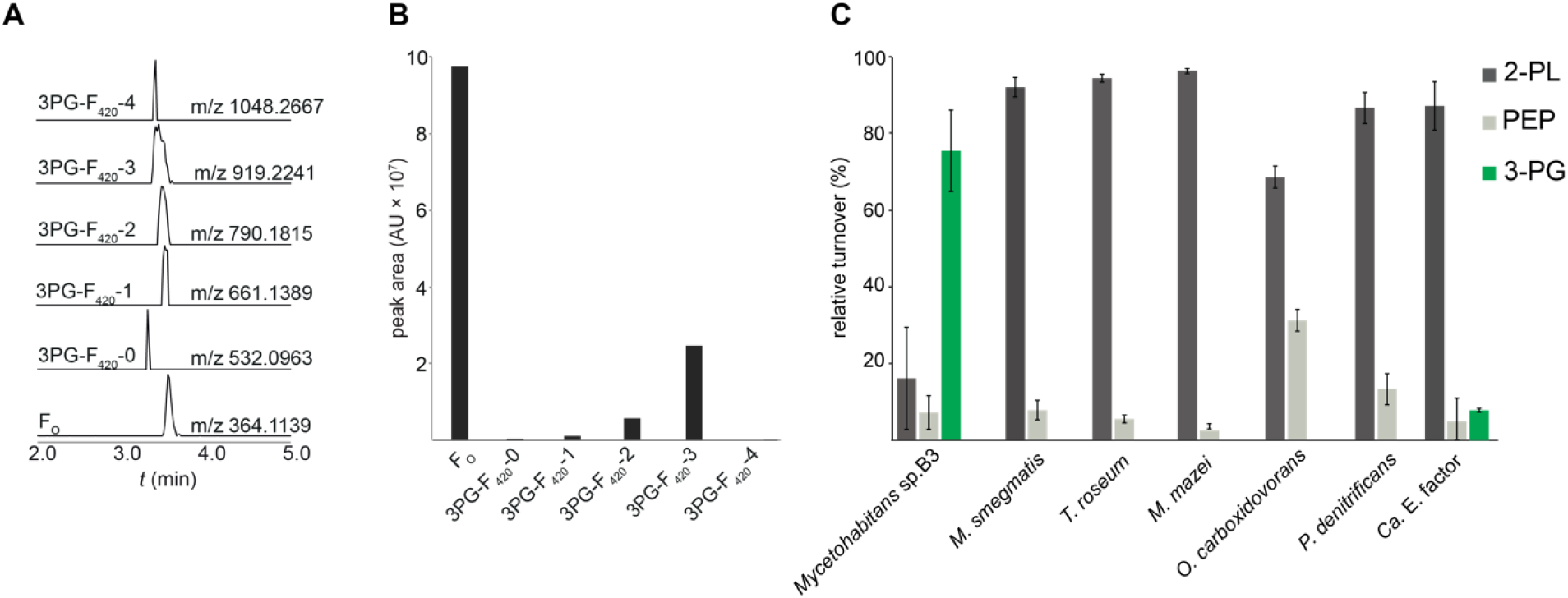
Production of 3PG-F_420_ and substrate specificity of CofC/FbiA enzymes. **A)** Extracted ion chromatograms (XICs) of deazaflavin species extracted from *Mycetohabitans sp*. B3 (peaks are scaled to the same height). **(B)** Areas under the peaks (arbitrary units) depicted in panel A. **(C)** Substrate specificity assay of CofC/FbiA from various source organisms. The CofC of *Mycetohabitans* sp. B3 showed strong 3-PG activation while all other homologs preferred 2-PL. *Mjan*-CofD was combined with all CofC homologs to perform the CofC/D assay. Error bars represent the standard deviation of three biological replicates. Relative turnover of 3-PG, 2-PL, and PEP are reflected by the rate of 3PG-F_420_-0, F_420_-0, and DF_420_-0 formation, respectively.

Next, we produced CofC of *Mycetohabitans* sp. B3 (*Myc*B3-CofC, accession number KQH55_09515) as a hexahistidine fusion-protein in *E. coli*, purified the enzyme by Ni-NTA affinity chromatography, and investigated its activity in a combined CofC/D assay using CofD from *Methanocaldococcus jannaschii* (*Mjan*-CofD). When the substrates were provided in equal concentrations in a competitive assay, *Myc*B3-CofC (**Figure 2C**) accepted 3-PG (65%), 2-PL (26.5%), and PEP (8.5%), a profile that was similar to the one obtained previously for the *Mrhiz*-CofC (22).

To obtain an overview of the substrate specificities of CofC we re-assessed CofC enzymes from well-studied F_420_ producing organisms such as *Mycolicibacterium smegmatis, Thermomicrobium roseum*, and *Methanosarcina mazei* (22) and assayed CofCs from further Gram-negative bacteria like *Paracoccus denitrificans, Oligotropha carboxidovorans* as well as the uncultivable *Candidatus* (*Ca*.) Entotheonella factor TSY1 that is rich in genes encoding F_420_-dependent enzymes (21). For all CofC-related enzymes analyzed, 2-PL was used most efficiently from all substrates compared. We observed PEP turnover in the range of 3.5% to 30% (**Figure 2C**). Generally, it can be concluded that CofC assays cannot discriminate whether 2-PL or PEP is the relevant substrate *in vivo*. The only CofC that accepted 3-PG to a certain extent (8%) was the enzyme from *Ca*. E. factor. However, compared to the *Mycetohabitans* enzyme there was no significant preference of 3-PG over PEP.

### Identification of 3-PG-binding residues of CofC

Next, we aligned primary amino acid sequences of CofC homologs to identify the residues that might be responsible for the altered substrate preference **(Figure 3A)**. A crystal structure of FbiD from *Mycobacterium tuberculosis* (*Mtb*-FbiD) in complex with PEP (PDB: 6BWH) showed eight amino acid residues to be in close contact with PEP suggesting a role in conferring the substrate specificity (32). Three of them are aspartate residues (D116, D188, D190) that complex two Mg^2+^ ions which in turn interact with the phosphate group of PEP. The remaining residues were supposed to bind the PEP molecule via side-chain atoms (K17, L92, S166) or backbone amino groups (T148 and G163). While most of these residues were highly conserved, two alignment positions showed a deviation in those residues able to activate 3-PG, namely L92 and G163 of *Mtb*-FbiD. While L92 is replaced by methionine (M91), the residue corresponding to FbiD-G163 was replaced by serine (S162) in *Mrhiz*-CofC and *Myc*B3-CofC. Homology modeling further suggested H145 to be a potential critical residue for 3-PG binding and C95 to be involved in the correct positioning of M91 **(Figure 3D, Figure S1**).

**Figure 3:**
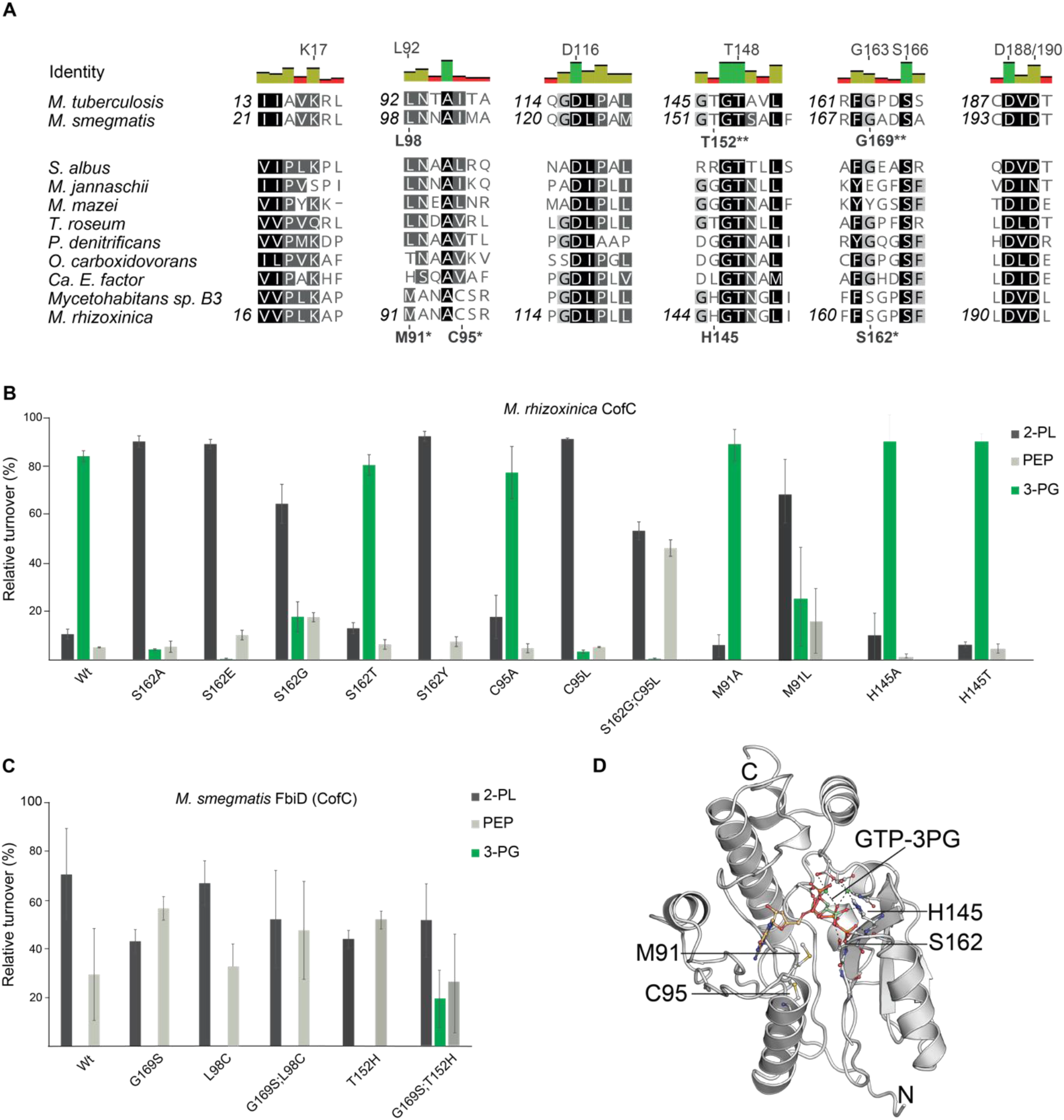
Residues determining substrate specificities of CofC homologs. **(A)** Multiple sequence alignment of CofC proteins from selected source organisms. Amino acids of *Mtb-*FbiD suggested previously to be involved in PEP binding residues (32) are indicated above the identity graph. Residues tested by mutagenesis are shown below sequences. Asterisks: crucial for 3-PG activation in *Mrhiz-*CofC, double asterisk: enabled 3-PG activation by *Msmeg*-FbiD. **(B)** Substrate specificity of *Mrhiz-*CofC after site-directed mutagenesis. Substitution of S162, C95, and M91 by residues occurring in 2-PL/PEP activating enzymes led to reduction or abolishment of 3-PG activation. **(C)** Substrate specificity of *Msmeg*-FbiD after site-directed mutagenesis. Gly169, Leu98, and Thr152 were exchanged by amino acids found in 3-PG activating homologs. While single mutations did not result in 3-PG activation, the combination G169S;T152H enabled 3-PG turnover. Error bars represent the standard deviation of three biological replicates. **(D)** Homology model of *Mrhiz*-CofC in complex with GTP (placed by molecular docking) and 3-PG (placed manually).

### Mutagenesis of CofC reveals S162 to be crucial for 3-PG activation

To probe the role of the suggested residues, we performed site-directed mutagenesis in *Mrhiz*-CofC (**Figure 3B**). Especially S162 turned out to be critical for 3-PG activation. While the least invasive mutation, S162T, retained most of the activity of wild-type CofC towards 3-PG, all other mutants of this residue preferentially turned over 2-PL and, to a lesser extent, PEP. This finding suggested that the hydroxy group present in S162 of WT and S162T might support the recruitment of 3-PG to the active site. M91L displayed reduced activation of 3-PG, while M91A was not impaired in 3-PG activation. Possibly, M91 controls the size of the substrate-binding pocket thus hindering (M91L) or facilitating (M91A) access of the larger substrate 3-PG to the active site. The C95A mutant approximately retained wild-type activity towards 3-PG, while C95L strongly reduced 3-PG activation. This was an indication that C95 might indeed affect the orientation of M91 and as a consequence 3-PG binding. Finally, the proposed interaction of 3-PG with H145 was not reflected in altered specificity profiles of H145A and H145T mutants of *Mrhiz*-CofC.

### Engineering *M. smegmatis* FbiD into a 3-PG activating enzyme

Inspired by the finding that S162 of the *Mrhiz-*CofC is necessary for 3-PG activation we wondered if mutation of the corresponding residue G169 of FbiD to serine (**Figure 3C**) could turn FbiD from *M. smegmatis* (*Msme*g-FbiD) into a 3-PG activating enzyme, thereby imitating the molecular processes underlying the evolution of 3PG-F_420_ biosynthesis. The G169S mutant, however, did not accept any 3-PG as substrate. We also tried to mutate L98 to facilitate the entry of 3-PG into the active site. However, neither the single mutant L98C, nor the double mutant G169S;L98C enabled 3-PG binding. Based on the homology model we suspected a residue corresponding to H145 of the *Mrhiz*-CofC might facilitate 3-PG binding. Indeed, while the single mutant T152H did not show any significant effect, the double mutant G169S;T152H successfully turned over 3-PG. The triple mutant (G169S;T152H;L98C) resulted in insoluble protein. Overall, these results showed that substrate specificity of *Msme*g-FbiD can be readily switched by changing only two residues and again supported a prime role of the critical serine residue for 3-PG recruitment.

### Structural insights into 3-PG activation by CofC

After several attempts had failed to crystallize recombinant *Mrhiz*-CofC we turned to *Myc*B3-CofC that was more soluble despite only minor differences in the amino acid sequence. From diffraction data collected to 2.4 Å the crystal structure could be solved by molecular replacement with a model of two superimposed structures (*Mtb-*FbiD and *Methanosarcina mazei* CofC, *Mmaz*-CofC). The overall structure was similar to the known homologs with the core of the single-domain protein being a six-stranded mixed β-sheet (**Figure 4**).

**Figure 4:**
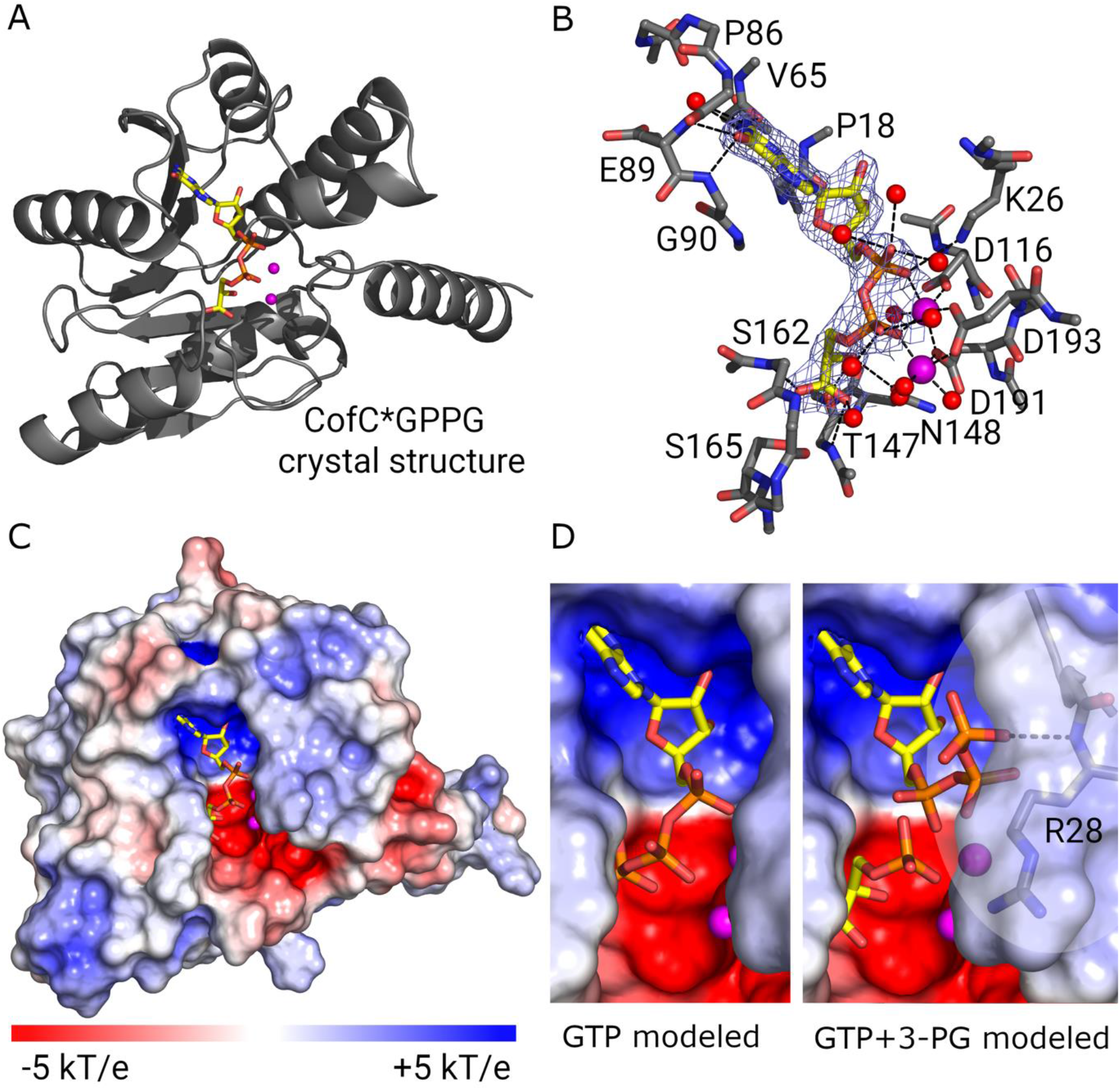
Structures of *Myc*B3*-*CofC in complex with GPPG (crystal structure) and educts (model). **(A)** Ribbon diagram of the crystal structure of CofC with its product GPPG. **(B)** Final electron density of GPPG (2F_o_F_c_ at 1.5 σ) and H-bonds for GPPG binding. Side chains are only shown for residues contacting the ligand. **(C)** Electrostatic surface representation of the protein shows a negatively charged deep pocket for GPPG. Two Mg^2+^-ions allow binding of the charged phosphates of GPPG. The electrostatic surface potential was calculated using APBS. Red: negative charge; blue: positive charge. **(D)** Possible educt conformations. GTP could position its α- and β-phosphates where GPPG binds to the Mg^2+^-ions (left side). With 3-PG binding in the same mode as the 3-PG moiety of GPPG the β- and γ-phosphates of GTP have to rotate out of the binding pocket, though (right side). The phosphate moiety of 3-PG is well poised for nucleophilic attack on the α-phosphate of GTP via a trigonal bipyramidal transition state. Figures were produced with PyMOL. A window around R28 is transparent to show the potential H-bond of GTP to the amide nitrogen. Electrostatic surface representations were made using the APBS electrostatics plugin in PyMOL. Carbon: yellow, oxygen: red, nitrogen: blue, magnesium: magenta.

Intriguingly, after initial refinement both molecules in the asymmetric unit independently showed unambiguous difference density for GPPG, the reaction product of GTP with 3-PG, completely immersed into the active site pocket. Each building block of GPPG is bound by several interactions (**Figure 4B, Table S1**). Guanine is distinguished from adenine by two H-bond donors to its oxygen (main-chain nitrogen of V65 and P86) and two H-bond acceptors to its amino group (main-chain carbonyl group of E89 and G90). The α- and β-phosphates are bound by two Mg^2+^ ions, which in turn are positioned by three aspartates (D116, D193, D191) similarly as had been shown for FbiD before (32). In the homologous structures the binding site for guanosine and ribose is almost completely conserved. Hence, GPPG is tightly bound, in fact, it turned out to be still quantitatively bound after purification of the protein from *E. coli* cell lysate, where both substrates were present.

The 3-PG moiety is primarily bound by H-bonds with the side chain of S165 to the carboxy group, apparently a highly conserved binding site of the carboxylate of the C3-acid as known from *Mtb*-FbiD (32). More interactions with the carboxy group arise from H-bonds with main-chain nitrogen atoms of T147 and S162, the latter being critical for selective 3-PG activation. Although the mild effect of S162T suggested otherwise, S162 did not interact via its hydroxy group with the ligand. Furthermore, there are H-bonds from the main-chain carbonyl groups of T147 and N148 to the 2-hydroxy group of 3-PG. Surprisingly, these binding partners of the 3-PG hydroxy group are structurally very well conserved and thus not likely to be involved in discrimination between 3-PG and 2-PL/PEP. The residue corresponding to *Mtb*-FbiD-K17 (*Mrhiz*-CofC-K20) did not interact with the substrate however, instead, K26 has taken over its role.

Taken together, the majority of the residues corresponding to the binding pocket residues of the PEP-binding FbiD (32) were also found to be involved in 3-PG binding. Although all C3-acids form the same hydrogen bonds, the carboxylate rotates by 36° about the axis through the carboxylate defined by the oxygens. This moves the phosphate by 2-3 Å. The hydrogen bonds for PEP have a more favorable geometry (average out-of-π-plane distortion 0.74 Å) than those for 3-PG (1.38 Å) but the phosphate group will not be positioned properly anymore to attack the α-phosphate of GTP. 3-PG with one more bond (carboxylate-C2-C3-O-P compared to carboxylate-C2-O-P in PEP or 2PL) can compensate for the new orientation of the carboxylate. The reason why the 3-PG adopts a new orientation is the main-chain rotation of S162, the CA and CB of which then squeeze the 3-PG into the productive conformation.

Based on the position of GPPG, the binding site for GTP is evident for the GMP moiety. Differential scanning fluorimetry (nano-DSF) measurements further corroborated the direct binding of GTP by CofC (K_D_ <20 µM) (see **Supplementary Methods, Figure S3**). The β-phosphate can either bind in the same position as the second phosphate of GPPG or point outward into the solvent close to R28N. The latter conformation allows the second substrate 3-PG to bind like PEP in FbiD. GTP and 3-PG are then positioned well for the reaction, the nucleophilic attack of the 3-PG phosphate on the α-phosphate of GTP (**Figure 4D**).

### Identification of further 3-PG accepting enzymes

After establishing that serine or threonine in the position corresponding to *Mrhiz-*CofC-S162 are linked to 3-PG formation, we hypothesized that the residue might be exploited as a diagnostic residue to identify further 3-PG activating enzymes. Going beyond highly related *Mycetohabitans* species, which can be expected to be 3PG-F_420_ producers, database searches revealed candidate proteins from as-yet uncultivated archaeal species **(Figure S2A)** that contained serine or threonine at the critical alignment position.

Since their source organisms were not accessible, we obtained the coding sequences of three of these candidate enzymes as synthetic genes and tested their substrate specificities **(Figure S2B)**. Interestingly, all of those enzymes accepted 3-PG as substrates. The circumstance that 2-PL was the best substrate of all three enzymes does not rule out the possibility that these enzymes are involved in 3PG-F_420_ formation given the fact that 2-PL is the default case for many enzymes examined in our assay system even if PEP is the natural substrate. Notably, two of the enzymes tested did not accept PEP as a substrate, a rather unusual finding. In the absence of 2-PL, this profile would result in the production of 3PG-F_420_. Taken together, S162 represents a diagnostic residue to predict the specificity of CofC towards 3-PG.

### Evolution of 3-PG accepting enzymes

To answer the question how 3-PG accepting enzymes might have evolved, we constructed a phylogenetic tree of CofC enzymes examined in this study (**Figure 5**). The *Mycetohabitans* CofC clade branched off early in the evolution of bacterial CofC enzymes and is neither closely related to nor derived from actinobacterial CofC/FbiD nor to other CofC enzymes found in Gram-negative bacteria. The archaeal 3-PG activating enzymes represent a monophyletic clade within the archaeal proteins. Taken together, we conclude that 3-PG activation has evolved independently in evolution at least twice from an ancestral 2-PL/PEP activating enzyme.

**Figure 5.**
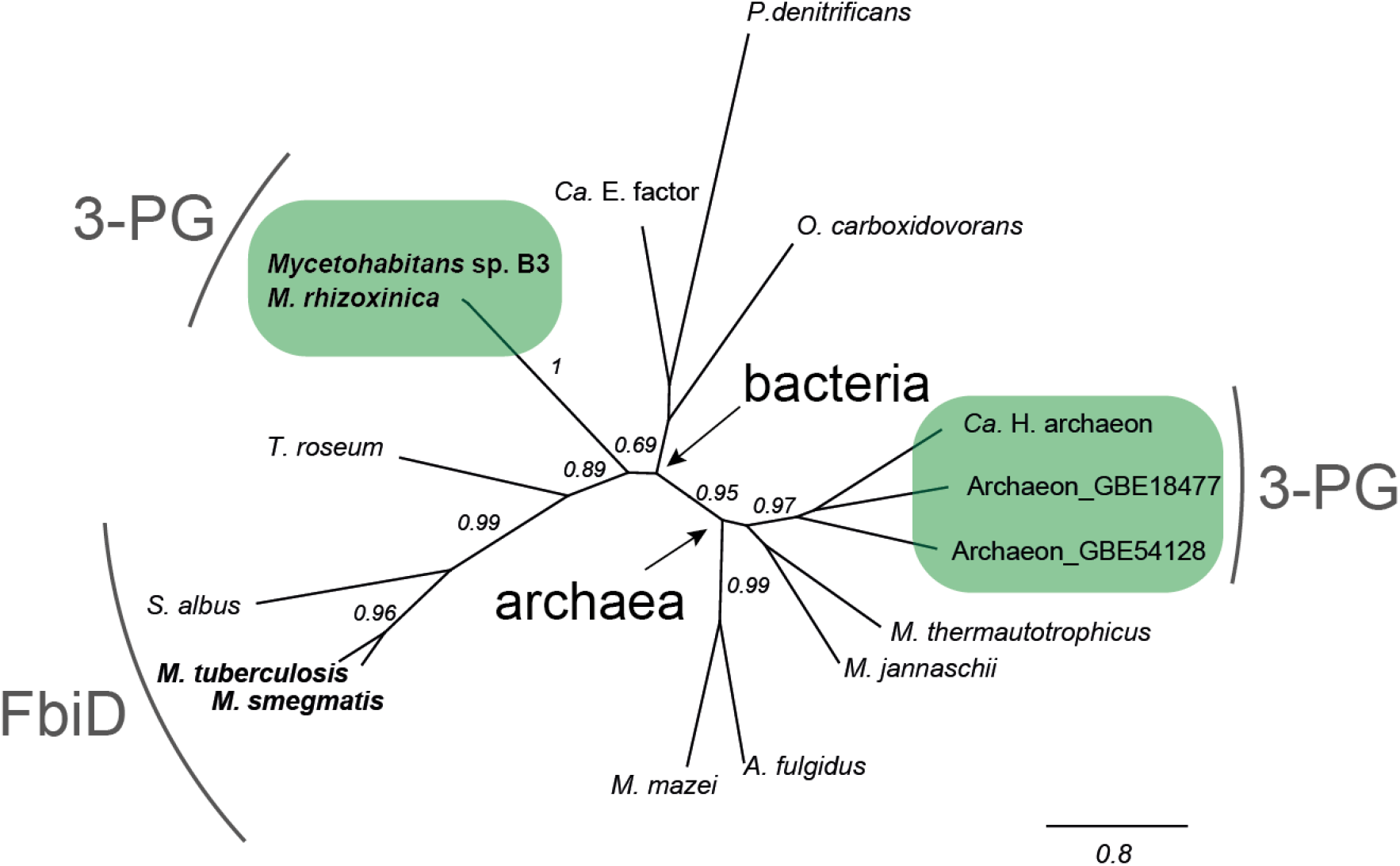
Phylogenetic tree of CofC/FbiD enzymes. The maximum likelihood method implemented in PhyML 3.0 was used to reconstruct the phylogeny. 3-PG activating enzymes (highlighted in green boxes) have evolved two time independently. Branch labels indicate SH-like support values. The scale bar represents substitutions per site. Accession numbers and source organisms of primary amino acid sequences are available in **Table S3**.

### Role of CofD in substrate specificity of F_420_ side-chain biosynthesis

After gaining insights into the substrate specificity of CofC a few questions remained. For instance, in almost all CofC homologs tested, 2-PL was the preferred substrate. This finding contrasted previous results obtained by Bashiri and colleagues (32) for *Mtb*-FbiD and CofC from *M. jannaschii* (*Mjan*-CofC), which was reported to accept exclusively PEP (32). Furthermore, the residual activity of *Mrhiz*-CofC and *Myc*B3-CofC towards PEP could not explain why PEP-derived DF_420_ was not found in their source organisms (22). We therefore assumed that the choice of the CofD homolog used in the combined CofC/D assay might have an impact on the overall outcome of the assay.

To test this hypothesis, we produced CofD homologs of several model species as hexahistidine-fusion proteins and performed CofC/D assays using several combinations of CofC and CofD. Strikingly, the choice of CofD homologs had a significant influence on the product spectrum of the CofC/D pair (**Figure 6**). For instance, when *Mrhiz*-CofC and *Myc*B3-CofC were combined with their cognate CofD, the apparent substrate specificity shifted almost completely towards 3-PG. The PEP-and 2-PL-derived products were only produced in traces by the CofC/D reaction. Similarly, when *Msmeg*-FbiD was combined with its natural partner *Msmeg*-FbiA instead of the homolog encoded in *M. jannaschii* (*Mjan*-CofD), the apparent activity towards 2-PL was almost entirely abolished. Obviously, CofD homologs have the ability to select between the pathway intermediates LPPG (2-PL-derived), EPPG (PEP-derived), and GPPG (3-PG-derived). In bacterial systems, CofD/FbiA appears to favor the intermediate that is known to be relevant in their source organisms, i.e., EPPG for Mycobacteria or GPPG for *M. rhizoxinica/Mycetohabitans* sp. B3. The archaeal *Mjan*-CofD, however, appears to prefer its natural substrate LPPG but displays relaxed specificity towards EPPG and LPPG.

**Figure 6:**
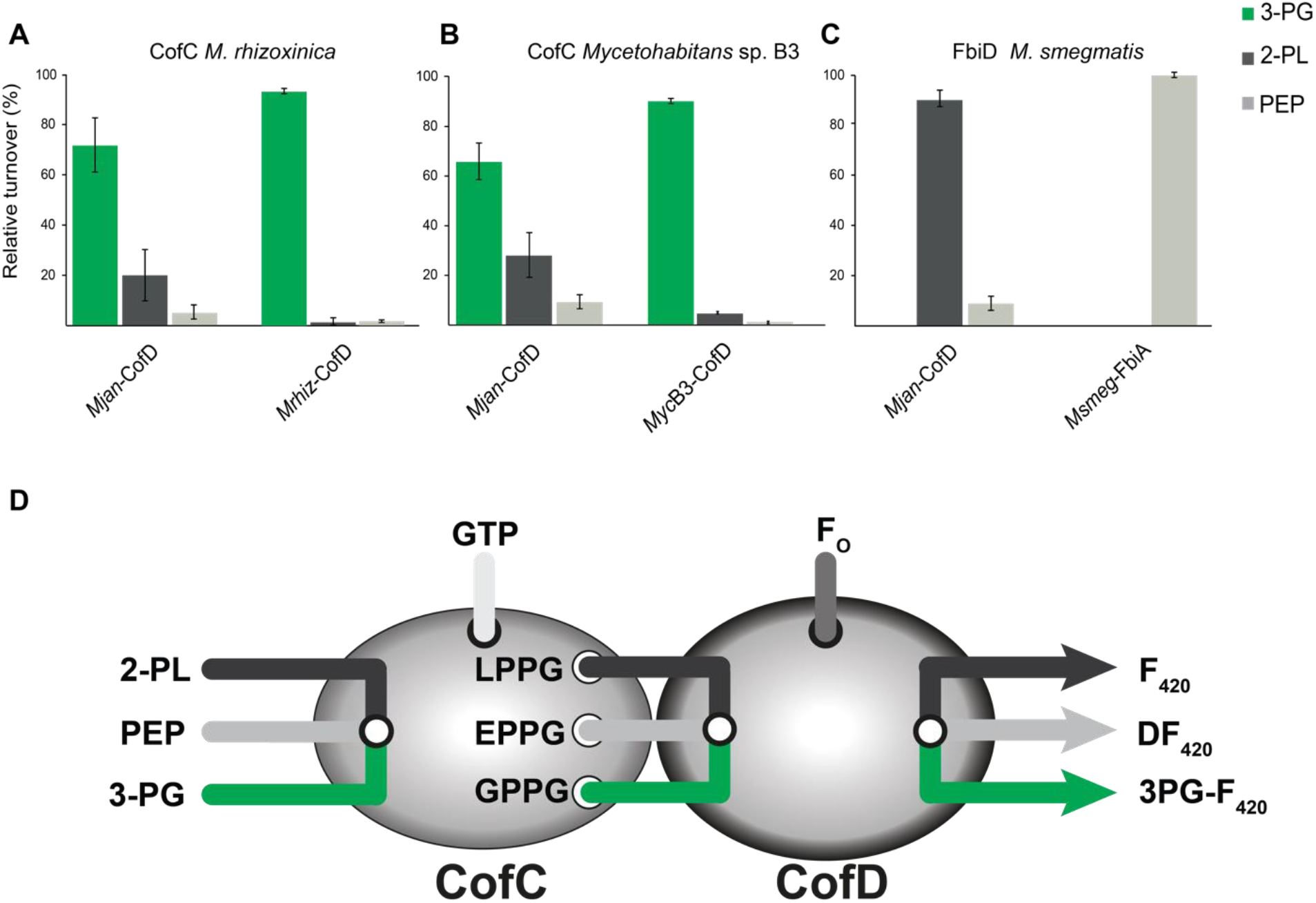
Influence of CofD/FbiA on the biosynthesis of F_420_-0 derivatives. **(A)** CofC from *M. rhizoxinica* **(B)** CofC from *Mycetohabitans* sp. B3, and **(C)** FbiD from *M. smegmatis* were tested with either *M. jannaschii* CofD (standard assay) or together with their cognate CofD enzyme from the same source organism **(D)** Schematic drawing illustrating the channeling of the labile metabolites LPPG, EPPG, GPPG from CofC to CofD. While CofC selects primary substrates, CofD is able to select between these intermediates. For abbreviations see Figure 1.

## Discussion

### Structural basis of CofC specificity

Extensive characterization of various CofC homologs, mutagenesis studies, and crystallography enabled us to spot residues responsible for the unusual substrate choice of CofC from *Mycetohabitans*. The crystal structure obtained from *Myc*B3-CofC revealed that most of the amino acid positions described for PEP-binding *Mtb*-FbiD play a role in 3-PG binding as well. However, rather than specific interactions with the free 2-hydroxy moiety, it is the conformation of S162 that forces the substrate into a position from which only the larger substrate 3-PG can undergo productive reaction of its phosphate group with GTP. The effect of M91 and C95 on substrate specificity shows that indirect influences on the overall conformation of the active site can be crucial for the correct positioning of the substrate. Nevertheless, the S162 residue proved as diagnostic residue to predict further 3-PG activating enzymes and even enabled engineering of *Msmeg*-FbiD into a 3-PG activating enzyme. Notably, a bidirectional change of the substrate specificity, i.e. from 3-PG to 2-PL/PEP in *Mrhiz*-CofC as well as from 2-PL/PEP towards 3-PG in *Msmeg*-FbiD FbiD as described here is a rather exceptional achievement.

### Evolution and occurrence of 3PG-F_420_ in nature

Interestingly, our mutagenesis study answers the question of how 3PG-F_420_ might have originated via mutation of 2-PL/PEP activating CofC on a molecular level. The phylogenetic tree suggests that 3-PG activating enzymes have evolved two times independently from an ancestral 2-PL/PEP activating CofC. Since DF_420_ is less stable than saturated forms, we suppose that the metabolic switching event has occurred to enable the formation of a stable F_420_-derivative in a metabolic background that lacked 2-PL or the DF_420_ reductase that is present in Actinobacteria (32) and Thermomicrobia (33). Here, we showed that the exchange of two amino acids in CofC/FbiD is mainly affecting substrate specificity and is thus sufficient to mimic this evolutionary process in the laboratory.

Considering that 3PG-F_420_ was detectable in biogas-producing sludge (22), there must be microorganisms outside the monophylogenetic clade of endofungal bacteria (*Mycetohabitans*) that produce 3PG-F_420_. The exploitation of the critical S162 in the active site indeed enabled us to identify further 3-PG activating enzymes, albeit from uncultured organisms. The source organisms may be related to the 3-PG producing organisms found in the digester. Efforts to isolate further 3PG-F_420_ producers remain ongoing and the knowledge gained here will facilitate this project in the future.

### CofD influences substrate specificity of CofC *in vivo* and *in vitro*

Another important key finding of this study is that CofC and CofD together contribute to the substrate specificity of the combined reaction, where CofD of some bacteria seems to represent a restrictive filter that acts after the more promiscuous CofC. This finding can explain the before-mentioned inconsistencies between results obtained *in vivo* and *in vitro*.

Even more importantly, we can now resolve the discrepancies concerning the results of CofC/D assays performed with CofC from archaea existing in the literature. Seminal work identified 2-PL to be a suitable substrate of archaeal CofC enzymes and thus proposed the original biosynthetic pathway to start from 2-PL (29). In contrast, a more recent study did not observe any turnover of 2-PL neither using the archaeal CofC from *M. jannaschii*, nor using FbiD from *M. tuberculosis* and suggested a biosynthetic route starting from PEP via EPPG, even for archaea (32). A follow-up study delivered further evidence supporting the biosynthetic route via DF_420_ in mycobacteria. The authors showed the formation of DF_420_ from PEP in cell-free extracts of *M. smegmatis* and could reveal strong binding of DF_420_ to the active site of FbiA (CofD) by X-ray crystallographic studies of the enzyme (34).

Our previous study (22), however, found 2-PL to be the best substrate of the *M. jannaschii* CofC enzyme - a result that we could reproduce here using CofC from *M. mazei*. It is also confusing that even the *M. smegmatis* FbiD tested in this study preferred 2-PL as substrate, again challenging the hypothesis that PEP is the preferred substrate of FbiD.

The solution to this perplexing situation comes with the herein defined influence of CofD/FbiA on the overall specificity of the CofC/D pair. CofD selects between the unstable pathway intermediates LPPG, EPPG, and GPPG (27). According to our results, *Msmeg*-FbiA exclusively accepted EPPG to form DF_420_. Bashiri *et al*. used the closely related *Mtb*-FbiA to perform CofC/D assays (32). Since *Mjan*-CofC can accept PEP as a minor substrate when combined with its cognate CofD (22), the assay resulted exclusively in DF_420_-formation when combined with FbiA, thus erroneously suggesting that 2-PL was not accepted by CofC. Similarly, the activation of 2-PL by FbiD remained “hidden” when assays were carried out with FbiA as a partner enzyme. Conversely, it is plausible that the “supernatural” turnover of 2-PL observed in all our assays performed with *Mjan*-CofD might be an artifact caused by the choice of a CofD homolog that might preferably turn over its natural substrate LPPG. For future studies towards the biosynthesis of novel F_420_ derivatives we, therefore, suggest that only a combination of CofC and its cognate CofD is suitable to reflect the *in-vivo* situation.

### The combined CofC and CofD reaction

The X-ray structure of *Myc*B3-CofC presented here included the reaction product GPPG, while the previous crystal structure of FbiD was obtained in the presence of PEP only (32). This is the first direct analytical evidence for the labile reaction product GPPG. So far, the existence of its congener LPPG was only confirmed by chemical synthesis followed by successful turnover by CofD (35). The fact that GPPG remains tightly bound to the enzyme points to a substate-channeling mechanism where the GPPG molecule is directly transferred to CofD to avoid degradation of the labile intermediate in the absence of F_O_. Substrate inhibition could also explain why any attempt to measure the activity of CofC in the absence of CofD remained unsuccessful (27,32) and why direct detection of GPPG or the related LPPG and EPPG from solution has failed so far.

The binding mode of GPPG also clearly revealed the GTP binding site of CofC. Notably, no evidence for GTP binding could be obtained experimentally for *Mtb*-FbiD and it was even speculated that GTP binding might require the presence of FbiA (32). This mechanism could be disproved for CofC of *M. rhizoxinia*.

## Conclusion

Taken together, this study represents a significant advance in understanding the flexibility of substrate specificity in CofC homologs and offers a molecular model for the evolution of 3PG-F_420_. By direct detection of the instable reaction product GPPG via X-ray crystallography we gained insights into the structural basis of the combined CofC/D reaction. The demonstration that CofC and CofD cooperate closely to control the entry of central carbon metabolites into the biosynthetic route to F_420_ derivatives also solved an ongoing debate in the literature and thereby re-established 2-PL as the most likely starting point of F_420_ biosynthesis in archaea. Future perspectives are opened up to investigate the cooperation of biosynthetic enzymes on a molecular level and to exploit this knowledge gained here for enhanced biotechnological production of coenzyme F_420_ and potentially novel derivatives.

## Materials and Methods

### Chemicals and microbial strains and metabolite extraction

Chemicals and media components were purchased from Acros Organics, Sigma Aldrich, Alfa Aesar, Carl Roth, and VWR. The *E. coli* strains BL21 (DE3), LOBSTR-BL21 (Kerafast), ETH101 and Top10 were grown routinely in Lysogeny Broth (LB). Axenic *Mycetohabitans* sp. B3 (NCBI genome accession JAHLKN000000000) was grown in 50 ml MGY media (22) at 30 °C at 110 rpm for one week. Cultures (OD_600_ = 2.5) were lyophilized, extracted with 10 mL ice-cold methanol, sonicated for 20 min and incubated at 250 rpm for 60 min. Cell debris was removed by centrifugation (4 °C, 10 min, 10000 rpm) and the supernatant was filtered into a round-bottom flask. The extract was dried in a vacuum rotary evaporator at 40 °C and re-dissolved in 1 mL LC-MS-grade water. Samples were analyzed using LC-MS as described before (22).

### Construction of expression vectors

Unless stated otherwise, primers (**Table S2A**) were designed using the software tool Geneious (36) and cloning was based on DNA recombination following the Fast Cloning protocol (37). The *E. coli* Top10 strain was used to propagate plasmids. PCR reactions were carried out using Q5 High-Fidelity polymerase (New England Biolabs) and oligomers used for amplifications listed in Table S1. Constructed plasmids (**Table S2B**) were confirmed by Sanger sequencing (Eurofins Genomics). CofC and CofD encoding plasmids (pMH04, 05, 10, 18, 19, 20,) were purchased from BioCat as codon-optimized synthetic gene construct cloned into pET28a+ between *Bam*HI and *Hind*III restriction sites. Plasmid sequences of pMH43 (pACYCDuet backbone), pMH59 (pET28), and pMH60 (pET28) were obtained by gene synthesis (BioCat), sequences are provided in the Supplemental Material **(Appendix)**.

### Site-directed mutagenesis

Putative substrate-binding residues of CofC were subjected to site-directed mutagenesis on DNA level using PCR (38). Amino acid numbering corresponds to the original residue position in the native CofC protein.

To mutate CofC from *M. rhizoxinica* (pFS03) at position 162 (S162), primer pairs oMH03/05, oMH05/06, oMH07/08, and oMH23/24 were used to generate S162E (resulting plasmid pMH22), S162Y (pMH23), S162G (pMH24), S162A (pMH25) and S162T (pMH26), respectively. Exchange of cysteine at position 95 (C95) was achieved using primer pairs oMH102/103 (C95L, pMH66) oMH104/105 (C95A, pMH67). The double-mutant C95L;S162G (plasmid pMH68) was obtained from pMH24 by amplification with primer pair oMH106/107. Mutation of histidine at position 145 either to alanine (H145A) or threonine (H145T) was obtained using primer pairs oMH116/117 (pMH74) or oMH118/119 (pMH75). Methionine at position 91 was substituted with alanine (M91A) or leucine (M91L) to yield pMH76 and pMH77.

*M. smegmatis cofC* (pMH10) served as a template for substituting glycine residue in position 169 (G169). Primer pairs (oMH33/34) or (oMH37/38) or (oMH45/46) or oMH47/48 were used to obtain G169S (pMH32), G169Y (pMH33), G169A (pMH34) or G169E (pMH35), respectively. Primer pairs oMH108/109 were used to substitute leucine (L98) with cysteine (L98C, pMH70). The double-mutants G169S; L98C and G169S;T152H were obtained from plasmid pMH32 using primer pair oMH110/111 and oMH124/125, yielding plasmid pMH69 and pMH80, respectively. The triple mutation was introduced in the pMH69 using oMH124/125 primers resulting in plasmid pMH81 (G169S;L98C;T152H).

### Heterologous protein production and purification

Production condition and purification of all *N*-terminal hexahistidine (N-His_6_) tagged proteins (CofC and CofD) were similar as described before (22). Accession numbers of native CofC/FbiD and CofD/FbiA proteins are listed in **Table S3**. In short, chemical competent *E. coli* BL21 (DE3) or LOBSTR-BL21 cells were transformed with individual CofC/CofD encoding plasmids and the respective antibiotic (kanamycin 50 µg/mL or chloramphenicol 25 µg/mL) was used to maintain selection pressure. Correct positive clones were grown overnight at 37 °C and 180 rpm and used to inoculate fresh 100 mL cultures (1:100). Upon reaching late exponential growth phase (OD_600_ = 0.7), expression of the gene was induced by the addition of 1 mM IPTG and incubated (18 °C, 180 rpm) following 18 hours for protein production. After harvesting, cells were disrupted with pulsed sonication. The clear cell lysate was loaded onto a Ni-NTA affinity column to separate N-His_6_ tagged protein. Later on, the protein was eluted with a higher concentration of imidazole (500 mM) and re-buffered in a PD-10 column.

### Combined CofC/D assay

Distinct derivatives of F_420_-0 were produced via biochemical reaction of purified CofC and CofD proteins in a combined assay (22,29). In-vitro reaction conditions were analogous to Braga *et al*. (22) and 50 µl reaction consisted of 100 mM HEPES buffer (pH 7.4), 2 mM GTP, 2 mM MgCl_2_, 0.14 nM Fo, 34 μM CofD, and 0.5 mM of substrates (3-phospho-D-glyceric acid, phosphoenolpyruvic acid, and 2-phospho-L-lactate). The reactions were initiated upon the addition of 26 μM CofC. Reactions were quenched with one volume of acetonitrile and formic acid (20%). Production of F_420_-0 derivatives was monitored in LC-MS. Technical set up, method, conditions for LC-MS analysis were similar as described before (22). Data analysis followed extraction of ion chromatograms (XICs), calculation of area under the curve (AUC), normalization of AUC, plotting area against time, and product formation was calculated for a linear time range (0 to 20 min). Quantification of relative product formation was determined from three biological replicates (n = 3) and plotted as bar charts. Standard deviations (SD) were used as error bars.

### CofC sequence alignment and phylogenetic tree inference

Multiple protein sequences of CofC from different F_420_ producing organisms (**Table S3**) were retrieved from the NCBI database and primary sequences were aligned based on their predicted structure using Expresso (T-Coffee) (39). For phylogenetic tree inference as implemented in Geneious Prime (36), the MUSCLE algorithm (40) was used to align sequences and a Maximum Likelihood tree was inferred using PhyML 3.0 (41) with the LG model for protein evolution and a gamma distribution of rates. Support values (Shimodaira-Hasegawa-like branch test) were computed and are shown above branches. Trees were visualized in Geneious Prime.

### Structural Modeling

The Phyre2.0 web portal was used to obtain a structural model for *M. rhizoxinica* CofC (PrCofC) (42). This enabled the identification of three enzymes (PDB: C3GX, PDB: 2I5E, PDB: 6BWH) that were used as a template for structural modeling resulting in models with 100% confidence in the fold. The enzyme FbiD from *M. tuberculosis* H37Rv (PDB: 6BWH) was used for further analyses. Initial structural alignment based on short fragment clustering of *M. rhizoxinica* CofC and FbiD was performed by program GESAMT (General Efficient Structural Alignment of Macromolecular Targets) from the CCP4i2 V1.0.2 program suite (43-45). This superimposed a total of 189 residues with an rmsd of 1.687 Å. FbiD binds to two Mg^2+^ ions that are important for catalysis and PEP binding (32). To place the Mg^2+^ ions in CofC, the CofC model was superposed on FbiD using residues surrounding the Mg^2+^ binding site. This model was used as a template for molecular docking of GTP into the CofC model using AutoDoc Vina. The input PDBQT files for AutoDoc Vina were generated with AutoDoc Tools V1.5.6 (46). The PRODRG server was used to generate the three-dimensional coordinates for GPPG from two-dimensional coordinates (47). The 3-PG was manually modeled in CofC•GTP using COOT (48). Representations of structures were prepared using PyMOL Molecular Graphics System (Schrödinger, LCC). The Adaptive Poisson-Boltzmann Solver (APBS) electrostatics plugin in PyMOL was used for the electrostatic surface representation (49).

### Crystallization and data collection

*Mycetohabitans* B3 CofC was further purified on a size exclusion column (Superdex75, 16/600, Cytiva) and concentrated in 50 mM Tris pH 7.4, 100 mM NaCl, 5 mM MgCl_2_, 2 mM mercaptoethanol (SEC buffer) to 8.2 mg/mL. Sitting drop crystallization trials were set up with screens Wizard I and II (Rigaku), PEG/Ion (Hampton Research), and JBScreen (Jena Bioscience) using 0.3 µL protein solution and 0.3 µL reservoir. Crystals appeared after 2 weeks with reservoir 10% PEG 3000, 200 mM MgCl_2_, 100 mM sodium cacodylate pH 6.5. After briefly soaking in a reservoir with 20% glucose added crystals were cryocooled in liquid nitrogen. Diffraction data were collected at BESSY, beamline 14.1. Data collection parameters are given in **Table S4**. Data sets were processed with XDSAPP (50).

### Structure solution and refinement

Programs used for this part were all used as provided by CCP4 (45). Sequence search against the PDB revealed structures of two related proteins, FbiD from *Mycobacterium tuberculosis* and CofC from *Methanosarcina mazei* with sequence identities of 30%. Molecular replacement by PHASER with 6BWG (*Mtb*-FbiD) or 2I5E (*Mmaz*-CofC) did not solve in the automatic mode, an assembly of both structures superimposed and truncated (residues 9-82, 91-173, 178-211 of monomer A of 6BWG and aligned residues of monomer A of 2IE5) solved the phase problem with an LLG of 233. The 6BWG structure was used as starting model for replacing the sequence with the *Mycetohabitans* sequence, one round of automatic model building with BUCANEER (51) and iterative rounds of refinement with REFMAC5 (52) and manual model building with COOT (53) completed the two protein chains.

Unambiguous water molecules were added when R_free_ reached 0.315 and a GDP moiety with two Mg^2+^ ions was built into the difference density in the active site. Pertaining difference density was connected to the β-phosphate suggesting a covalently bound PEP or 3-PG. Only the latter refined without residual difference density above ±2σ. For final refinement, non-crystallographic symmetry was not used. TLS refinement was applied with one group per monomer. Refinement statistics are given in **Table S4**.

The C3-acid substrates are fixed in the active site by three H-bonds of their carboxylate group. For H-bonds to π-systems (peptide bond, carboxylate group) the partner should lie in the same plane. To characterize deviations from favorable H-bonding geometry we calculated the average distance of the partner to the π-plane.

### Data deposition

The crystal structure of CofC from *Mycetohabitans* sp. B3 was deposited at the protein database PDB (PDB code: 7P97).

## Author contributions

MH, DB (cloning, CofC/D assays, mutagenesis, LC-MS analysis), and DL (diagnostic sequence analysis) performed research and analyzed data, and contributed to writing the manuscript. IR performed research (cultivation and metabolite extraction). SS and LB performed research (crystallization, GTP binding assays). GJP and ML analyzed data (X-ray crystallography, structural modeling, GTP binding assays) and contributed to writing the manuscript. GL performed research (phylogenetic reconstruction), designed the study together with ML, and wrote the manuscript.

## Acknowledgments

GL, MH and DB thank the Carl Zeiss Foundation, GL and DL the German Research Foundation (DFG; Deutsche Forschungsgemeinschaft) Grant LA4424-1/1 (Project: 408113938) and the Leibniz Association for funding. ML and SS thank the DFG Grant LA2984-5/1 (Project: 389564084) and LA2984-6/1 (Project: 449703098) for funding. We declare there are no conflicts of interest. IR is grateful for financial support from the European Union’s Horizon 2020 Research and Innovation Programme under the Marie Skłodowska-Curie grant agreement No. 794343.

